# Pervasive head-to-tail insertions of DNA templates mask desired CRISPR/Cas9-mediated genome editing events

**DOI:** 10.1101/570739

**Authors:** Boris V. Skryabin, Leonid Gubar, Birte Seeger, Helena Kaiser, Anja Stegemann, Johannes Roth, Sven G. Meuth, Hermann Pavenstädt, Joanna Sherwood, Thomas Pap, Roland Wedlich-Söldner, Cord Sunderkötter, Yuri B. Schwartz, Juergen Brosius, Timofey S. Rozhdestvensky

## Abstract

CRISPR/Cas9 mediated homology-directed DNA repair is the method of choice for precise gene editing in a wide range of model organisms, including mouse and human. Broad use by the biomedical community refined the method, making it more efficient and sequence specific. Nevertheless, the rapidly evolving technique still contains pitfalls. During the generation of six different conditional knock-out mouse models, we discovered that frequently (sometimes solely) homology-directed repair and/or non-homologous end-joining mechanisms caused multiple unwanted head-to-tail insertions of donor DNA templates. Disturbingly, conventionally applied PCR analysis—in most cases—failed to identify such multiple integration events, which led to a high rate of falsely claimed precisely edited alleles. We caution that comprehensive analysis of modified alleles is essential, and offer practical solutions to correctly identify precisely edited chromosomes.

## Introduction

Genome editing is a powerful research tool for biology and medicine. In recent years, considerable progress has been made in this area as a result of new technologies that have emerged to directly modify genes at the stage of single-cell embryos (zygote), stem cells, including iPSC, or germ cells. Discoveries and application of sequence-specific programmable nucleases: 1) ZFN (Zink-Finger Nucleases) ^1^, 2) TALEN (Transcription Activator-Like Effector Nucleases) ^2^, and 3) CRISPR/Cas9 ribonucleoprotein complexes constitute some of the advances ^3, 4^. CRISPR (Clustered Regularly Interspaced Short Palindromic Repeats) are prokaryotic genomic short palindromic repeats located in clusters ^5^. These clusters are transcribed and processed into small RNAs interacting with Cas9 protein resulting in a sequence-specific endonuclease ^6^. The CRISPR/Cas9 complex comprises of two RNA molecules: crRNA (CRISPR RNA) and tracrRNA (trans-activator for crRNA) ^7^. The crRNA contains ∼20 nucleotides of recognition sequence complementary to the targeting region of DNA, whereas tracrRNA interacts with Cas9 protein and base-pairs with a crRNA ^8^. The minimal “artificial” CRISPR/Cas9 complex consists of a crRNA-tracrRNA molecule hybrid (guide RNA or gRNA) and Cas9 protein-DNA endonuclease ^9^. Cas9 is a 1368 aa multi-domain protein isolated from *Streptococcus pyogenes* (SpCas9); that together with the crRNA-tracrRNA complex cleaves double stranded DNA (dsDNA) adjusted to the PAM-motif (protospacer adjacent motif; NGG-sequence) within the DNA strand complementary to crRNA (target strand) using the HNH-like nuclease domain and the opposite, non-target strand, via the RuvC-like domain ^10^. The CRISPR/Cas9 complex has been broadly used to generate defined site-specific cleavage of genomic DNA; it is a fast, inexpensive and effective DNA editing system that has a wide range of potential applications. In living cells, the sequence-specific dsDNA breaks are repaired by non-homologous end joining (NHEJ) or homology-directed repair (HDR) mechanisms. NHEJ often results in small insertions or deletions at the dsDNA break site, which may impair the function of a targeted gene. The NHEJ mechanism is commonly utilized for the generation of conventional gene knock-out models in a wide range of organisms. The HDR mechanism requires a specific donor DNA template, most often co-injected together with the CRISPR/Cas9 complex, and results in precise genome editing events. HDR enables the insertion of specific point mutations, the addition of in-frame translated epitopes, the performance of sequence-specific knock-in (KI) events of genes, the generation of conditional knock-out (cKO) genetic models, etc.. Once refined to perfection, CRISPR/Cas9-mediated HDR-based genome editing holds immense promise for gene therapy. Indeed, much of the genome-editing community was invested in improving the efficiency and sequence specificity of the CRISPR/Cas9 complexes ^11–19^. However, several limitations of the technique, such as low efficiency of HDR, off-target effects or genomic rearrangements remain challenging obstacles ^20, 21^.

Our study examines the generation of six conditional knock-out mouse models that employed CRISPR/Cas9 mediated homology-directed repair mechanism in ten knock-in procedures. A comprehensive analysis revealed that direct genome editing of zygotes had resulted in mosaic genotypes of targeted mice (F0 generation). Surprisingly, more than half of the F1 offspring with modified loci displayed multiple head-to-tail donor DNA integrations. We demonstrated that both homology-direct repair (HDR) and/or non-homologous end-joining (NHEJ) mechanisms were utilized. Importantly, conventionally applied PCR analyses using the outside targeting homology flanking primers, erroneously displayed integration of the desired single copy template; thus, the analysis failed to identify insert multiplication. If undetected, this would undermine the validity of studies involving such animal models. To avoid this shortcoming, we suggest methods that improve analyses and verification of correctly targeted loci.

## Results

### Generation and analysis of F0 founders for conditional knockout mouse models

The strategy to generate conditional KO mouse models by simultaneous CRISPR/Cas9 mediated insertions of two *LoxP* sites using two crRNA and two single-stranded oligonucleotides (ssODN) (2sgRNA-2ssODN), proposed by Yang et. al. ^22^ has been shown to be inefficient in an extensive study involving more than 50 different genomic loci ^23^. Our alternative “one-step” strategy for the generation of conditional KO mouse models using CRISPR/Cas9 complexes and long donor DNA templates, containing two *LoxP* sites is similar to those recently reported ^24, 25^, and could be exemplified by *S100a8* (calcium-binding protein A8) gene targeting. Based on computational analysis, we predicted that genomic elimination of the second exon would result in a translational frame shift leading to *S100a8* gene inactivation. Therefore, we designed a donor DNA fragment with *LoxP* sites flanking the second exon of *S100a8* gene (Fig. 1A, 2A-D). Our general strategy for one-step insertion of both *LoxP* sites relied on the active cellular HDR mechanism. We constructed a DNA template harboring exon-intronic regions flanked by *LoxP* sites with relatively short (77/83 nt) PAM-mutated homologous arms (Fig. 1A, 2B). In order to select CRISPR/Cas9 complexes that efficiently cut genomic DNA at a chosen position, we designed at least three sequence-specific crRNAs for each flanking region. To gauge whether selected crRNA pairs efficiently guide genomic deletion *in vivo*, we injected Cas9 components with different combinations of crRNAs into fertilized mouse oocytes. Subsequent PCR amplification of loci between pairs of crRNAs determined the efficiency of CRISPR/Cas9 complex targeting (Fig. S1). The most efficient crRNA pair together with the donor DNA template and Cas9 components were then microinjected into the cytoplasm of fertilized mouse oocytes (Table S1). For the *S100a8* project, we obtained 34 pups (F0 generation) from 193 modified embryos. Initially, the selection of positively targeted mice was performed by PCR amplification of the genomic DNA region with d3 and r3 primers located outside the donor DNA flanking homology region (Fig. 1A, B). We detected appropriate (∼700bp) PCR products representing a potentially desired targeted locus for mouse number 11 only (Fig. 1B). The other animals contained either wild type (∼550 bp) or deletions surrounding the targeted *S100a8* exon-intronic region (Fig. 1B). The infrequent HDR events in combination with negative amplification results for most of the animals prompted us to investigate all mice with a different PCR approach; thus, we decided to amplify sequences adjacent to the *LoxP* sites paired with primers located in the corresponding genomic flanks. We used PCR primers d3/Ar1 and Ad1/r3 for the 5’- and 3’- regions, respectively as shown in Fig. 1A, C and D. Founder (F0) number 11 was confirmed to contain the correctly targeted allele, but an additional founder (number 6) was positively identified (Fig. 1C, D). Notably, six mice that were previously identified as harboring only wild type alleles or deletions within the targeted region revealed the presence of at least one potentially HDR integrated *LoxP* site. These observations pointed to a mosaicism of F0 founders. To exclude false positive PCR identification of founder number 6, we performed gradient PCR amplification of donor DNA together with flanking regions using a combination of either d4/r4 or d4/r3 primers (Fig. 1A, E, Fig. S2). In both amplification schemes, only a single PCR product was detected—indicating correct HDR integration of a single copy donor DNA template (Fig. 1E, Fig. S2).

**Figure 1.**
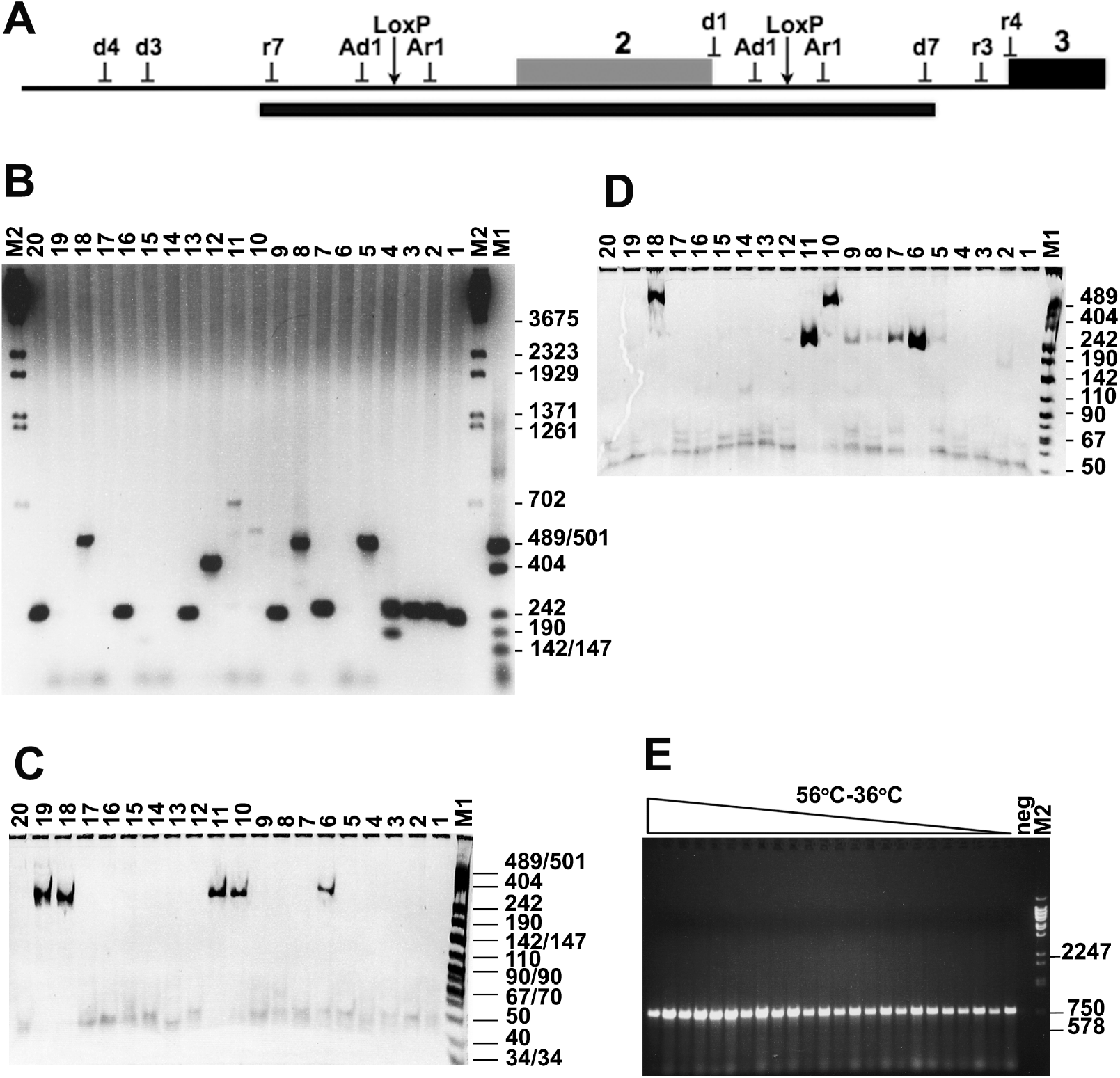
PCR analysis of the *S100A8* targeted locus. (**A**) Genomic structure of the targeted locus with positions of PCR primers (d1, d3, d4, d7, r3, r4, r7), where (d) denotes direct and (r) denotes reverse orientation. Intronic and intergenic regions are shown as line, exons are shown as filled boxes numbered above. The oligonucleotide pairs Ad1 and Ar1 are not present in the mouse genome but introduced as diagnostic sequences together with the LoxP sites. The black bar below corresponds to the location of the DNA template employed. (**B**) PCR analysis of genomic DNA from F0 founder mice 1-20 (labeled above), using primer pair d3/r3 located outside of the DNA template homology arms (Figure 1D). The PCR products of 715 bp and 543 bp correspond to the correctly targeted and wild type alleles of *S100A8* gene, respectively. The PCR products (>715 bp) originating from multiple head-to-tail integrations of the DNA template were not detected in the founder analyzed. Size marker positions (in bp) are shown on the right. (**C, D**) PCR analyses of DNA samples from F0 founder mice 1-20 using primer pairs d3/Ar1 (**C**) and Ad1/r3 (**D**). (**C**) The PCR product of 257 bp corresponds to HDR integration detected in mouse samples 6, 10, 11, 18 and 19. (**D**) The expected PCR product of 204 bp was detected in animals 6-9 and 11. (**E**) PCR analysis at different annealing temperatures of genomic DNA from F0 founder number 6 using primer pair d4/r3. Only one PCR product (750 bp) corresponding to a single copy targeted locus was detected.

**Figure 2.**
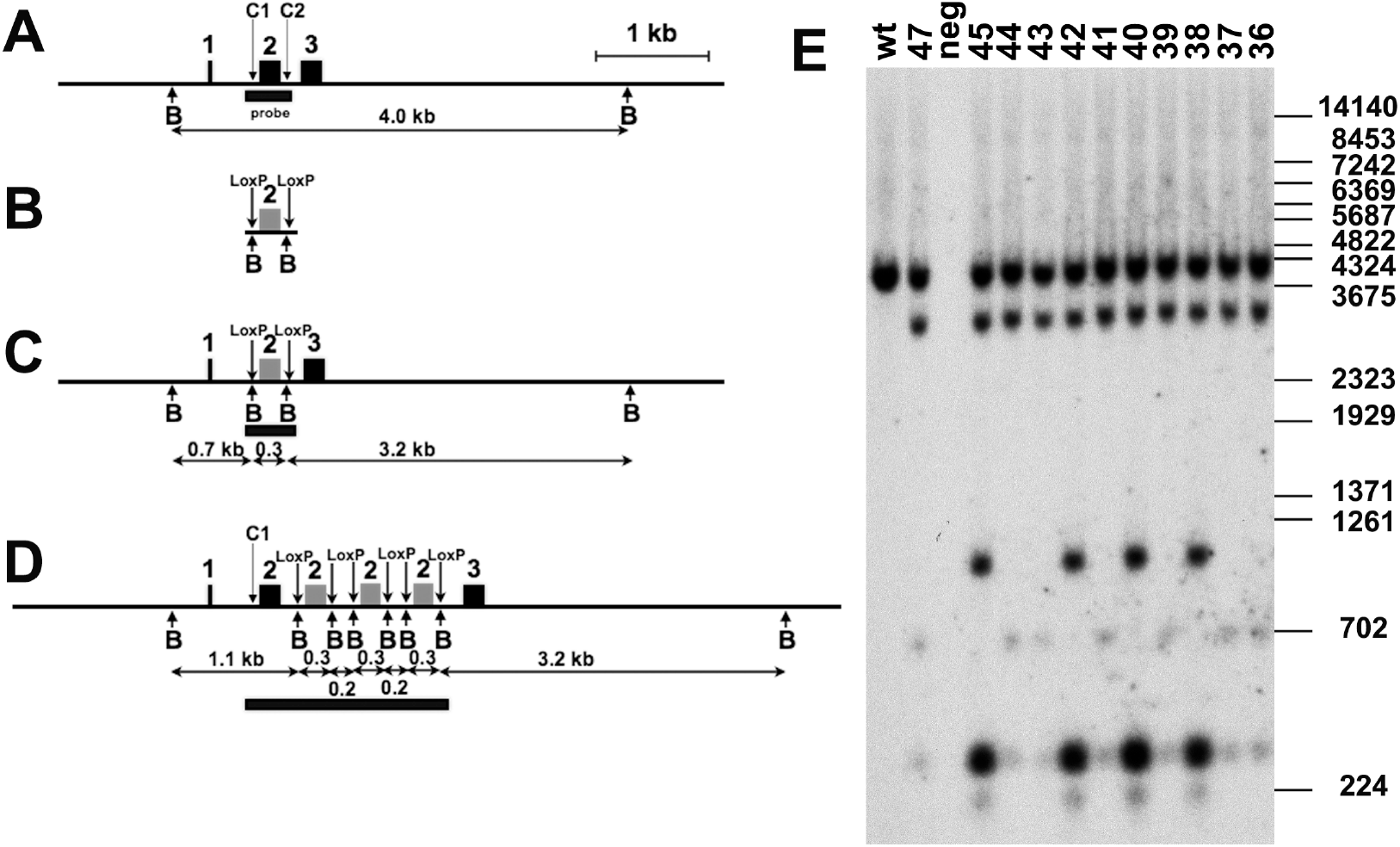
Schematic representation of *S100A8* gene targeting strategy. Exon 2 was chosen for elimination. Intronic and intergenic regions are shown as line, exons are shown as filled boxes numbered above. The vertical arrows indicate the target sites for the CRISPR/Cas9 complex with crRNA3 (C1) and crRNA12 (C2), respectively. The arrows marked with “B” correspond to *Bam*HI restriction endonuclease sites. The black bars below (marked “probe” in A) correspond to areas recognized by donor DNA specific probes used in Southern blot analyses. The horizontal arrows denote the expected sizes of restriction DNA fragments given in kb. (**A**) Wild type locus. (**B**) Donor DNA template used in this study; the two LoxP sites are indicated by vertical arrows. (**C**) Genomic locus after HDR with single copy integration. (**D**) Targeted genomic locus with triple insertion of the donor DNA template. (**E**) Southern blot analysis of genomic DNA of the F1 offspring (36-45, 47) hybridized with the template-specific probe (labeled in A). *Bam*HI enzymatic digestion revealed the wild-type allele (4.0 kb) and three DNA fragments (3.2, 0.7 and 0.3 kb) corresponding to the targeted allele (marked in C). DNA samples 36, 37, 39, 41, 43, 44, and 47, contain the correctly targeted *S100A8* allele (*S100A8 ^+/-^*). Samples 38, 40, 42, and 45 contain DNA fragments of 1.1kb and 0,2 kb in size, indicating multiple copy head-to-tail integrations at the targeted locus (marked in D). Size marker positions (in bp) are shown on the right. The DNA sample from the wild-type control mouse is indicated as “wt”.

### Analysis of F1 generation mice revealed mosaicism of F0 founders

The offspring obtained after crossing of *S100a8* conditional KO founder number 6 with wild type mice was further analyzed by PCR and sequencing. Surprisingly, we detected two types of locus targeting. In the first, we confirmed the correct *CRISPR*/*Cas9* nuclease *C1* and *C2* cleavage of the genomic DNA locus and single copy integration of donor DNA template via HDR mechanism in offspring numbers 36, 37, 39, 41, 43, 44 and 47 (Fig. 2C, E). The correct integration at the nucleotide level was confirmed by sequencing. The second type corresponded to tandemly multiplied DNA template integration yielding up to 3 copies (confirmed by Southern blot, quantitative PCR (qPCR), digital droplet PCR (ddPCR) analyses) in a head-to-tail configuration at a single *CRISPR*/*Cas9* nuclease *C2* mediated DNA break (Fig. 2D, E, Fig. 3, Fig. S3). The 3’-end of the DNA fragment integrated via HDR, while the 5’-end integrated via a non-homologous end-joining (NHEJ) mechanism (Fig. 3. Fig. S3). As discussed in detail below, head-to-tail multiplications of donor DNA were obtained for other eight KI projects involving six different gene loci (Table 1).

**Figure 3.**
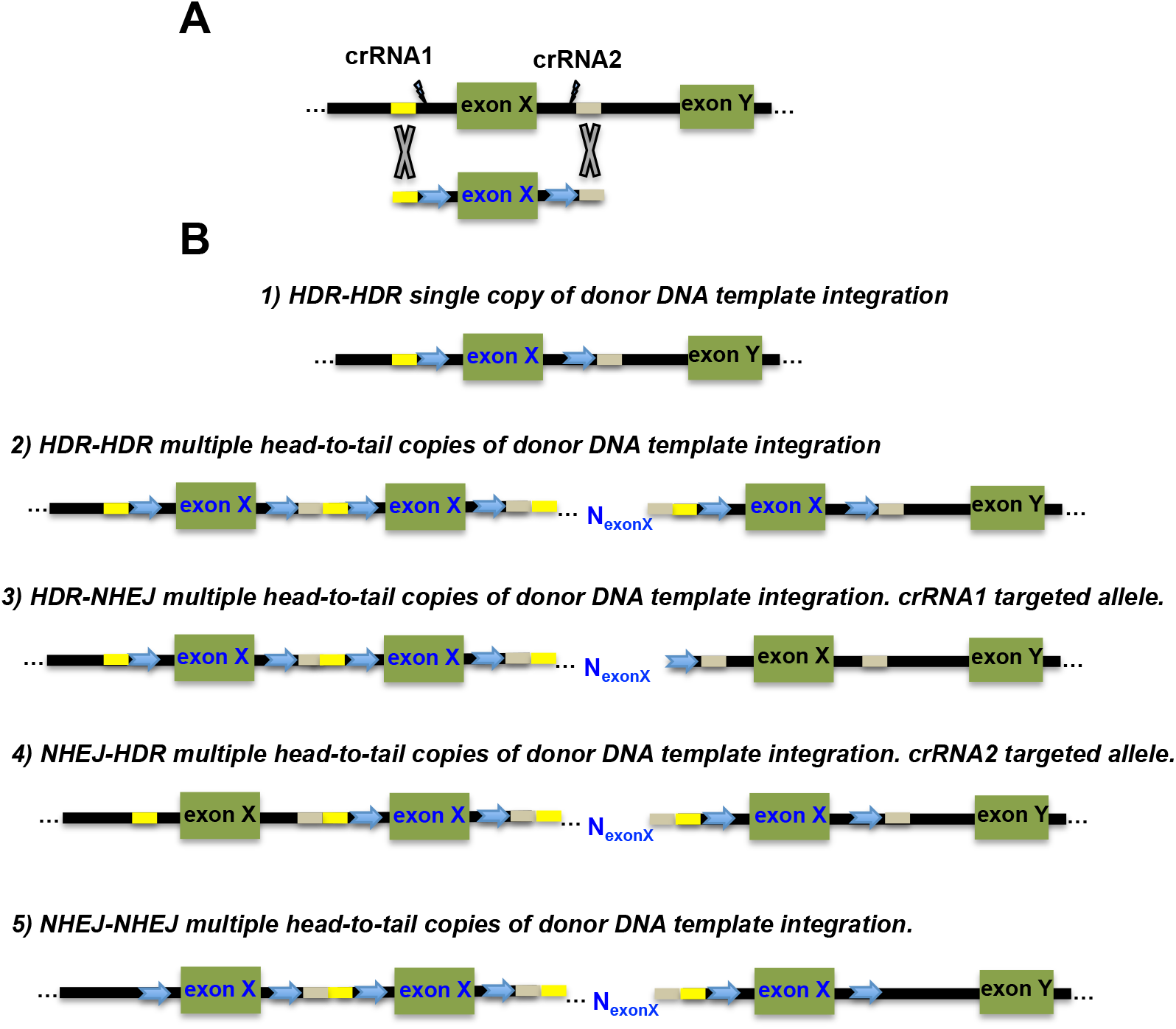
(A) Schematic representation of the loci for the conditional KO targeting strategy. Intronic regions are shown as lines, exons are shown as filled boxes. Homologous arms for HDR are marked in yellow and grey for left and right flanks, respectively. The target sites of the CRISPR/Cas9 complex are denoted as crRNA1 and crRNA2, respectively. (**B**) Different types of donor DNA integrations.

**Table 1.** Summary of conditional KO loci targeting and mechanisms of donor DNA integrations. *Gene name*: the names for conditional KO targeted genes are indicated (official ID provided by MGI). *Nr. of F0 selected animals*: number of F0 founders selected to contain a positively targeted allele. *Nr. of F1 analysed animals*: number of analysed mice from the F1 generation. *Nr. of F1 positive SC animals*: number of mice with correct HDR-HDR single copy donor template integration. *Nr. of F1 positive MC animals*: number of mice with identified multiple integrated copies of donor template. *Template size/strandedness (ss-ds DNA)*: donor DNA template sizes and strandedness are indicated. *Mechanism*: mechanism for donor DNA template integration as determined.

| Gene name | Nr. of F0 selected animals | Nr. of F1 analysed animals | Nr. of F1 positive SC animals | Nr. of F1 positive MC animals | Template size/strandedness (ss-ds DNA) | Mechanism |
| --- | --- | --- | --- | --- | --- | --- |
| *S100a8* | 2 | 14 (**nr.6**)<br>7 (**nr.11**) | 9 (**nr.6**)<br>4 (**nr.11**) | 5<br>3 | ssDNA (PCR)<br>**591 nt** | HDR-HDR<br>NHEJ-HDR<br>NHEJ- NHEJ |
| *Trek1* | 2 | 21 | 0 | 16* | dsDNA<br>**1257 bp** | HDR – NHEJ<br>NHEJ-HDR |
| *Trek1* | 6 | 28 | 0 | 12 | ssDNA (PCR)<br>**1257 nt** | HDR-HDR<br>NHEJ-HDR<br>HDR - NHEJ |
| *Trek1* | 3 | 26 | 0 | 3(F0) | ssDNA (IDT)<br>**1286 nt** | nd |
| *Inf2* | 11 | 34 | 2 | 15 | ssDNA (PCR)<br>**711 nt** | HDR-HDR |
| *Trpc6* | 3 | 22 | 5 | nd | dsDNA<br>**880 bp** | HDR-HDR |
| *Trpc6* | 4 | 34 | 5 | 2 | ssDNA (PCR)<br>**880 nt** | HDR-HDR |
| *Ccnd2* | 1 | 46 | 19 | 1(F0) | dsDNA<br>**1658 bp** | HDR-HDR |
| *Il4_5'LoxP* | 18 | 49 | 19 | 30 | ssDNA (PCR)<br>**210 bp** | HDR-HDR |
| *Il4_flox* | 4 | 41 | 0 | 1(F0) | ssDNA (PCR)<br>**1258 nt** | NHEJ-HDR |
|  |  |  | Total F1<br>63<br>(~43%) | Total F1<br>83<br>(~57%) |  |  |

### PCR analysis of animals with multiple head-to-tail DNA template integration

As previously mentioned, PCR analysis of F0 animals using primers flanking homologous arms of DNA inserts did not reveal the presence of multiple tandem duplications in the targeted locus at various PCR amplification parameters; this includes different primers, as well as various touchdown and annealing temperatures (Fig. 1E, Fig. S2). For all other “one-step” conditional KO projects, we only detected amplification products indicating “single copy insertion” (Fig. S4C).

Considering difficulties in identifying head-to-tail insertions when relatively long donor templates were used (from 550 bp to 1,65 kb), we tested the HDR mediated integration of a single stranded DNA (ssDNA) harbouring one *LoxP* site (∼210 nt) during the construction of an *Il4* gene conditional mouse model (Fig. 4A). Multiple head-to-tail integrations of a single *LoxP* site were verified in the F1 mouse offspring. Altogether, 49 mice were PCR analysed using primers (SD1 and SR1) flanking the *LoxP* site homologous arms (Fig. 4A). Tandem multiplication of the *LoxP* harbouring DNA template was detected in 5 mice: numbers 34, 40, 42, 44, 48, all other mice revealed a PCR product corresponding to a single copy *LoxP* integration into the *Il4* gene locus using the HDR-HDR mechanism (Fig. 4B). This relatively low frequency of head-to-tail amplification was suspicious. Hence, we developed and performed additional control PCR amplification by using non-overlapping bidirectional primers (SD1r and SR1d) that would specifically detect head-to-tail *LoxP* repeats (Fig. 4A). Surprisingly, a total of 30 mice containing multiple copies of donor DNA were detected, indicating that ∼83% of mice harbouring *LoxP* head-to-tail multiplications were not verified by standard, commonly used PCR detection methods (Fig. 4C).

**Figure 4.**
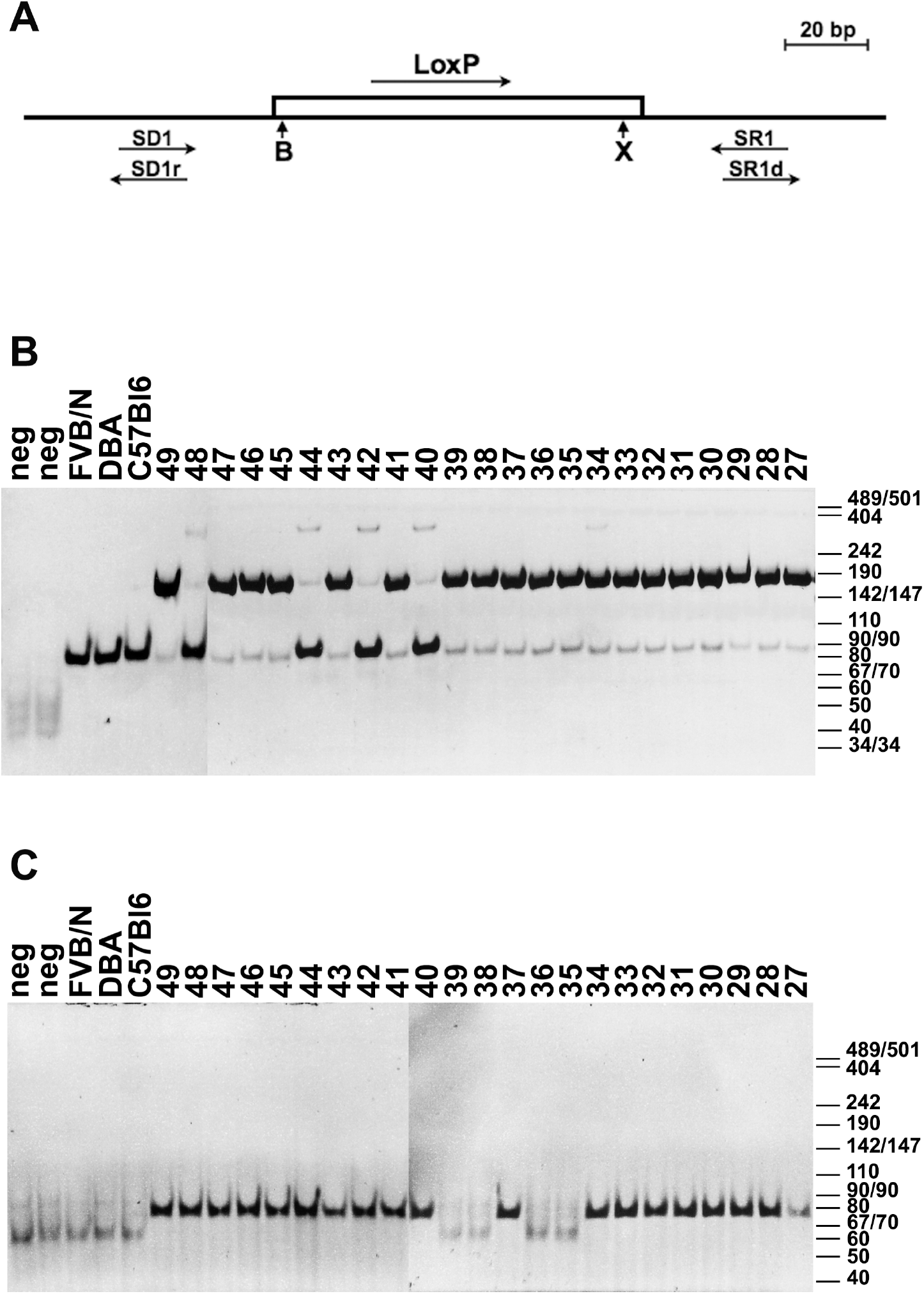
Analysis of F1 mice for *LoxP* site integration in the *Il4* locus. (**A**) Schematic representation of the *IL4* 5’-*LoxP* DNA template. The genomic region is shown as line, the inserted artificial DNA sequence is indicated as open rectangular. The 5’-*LoxP* site is designated with an arrow above, the restriction endonuclease sites *Bam*HI (B) and *Xho*I (X) are indicated below. PCR primers are denoted by arrows. (**B**) PCR analysis of genomic DNA from selected F1 founder mice using the SD1/SR1 primer pair. (**C**) PCR analysis of genomic DNA from selected F1 founder mice using bidirectional primers SD1r and SDR1d specifically detecting head-to-tail *LoxP* target DNA repeats.

### Southern blot analysis of targeted genomic loci

Alerted by the high false-positive rate of conventional PCR analysis, we turned to Southern blot hybridization to test whether multiple head-to-tail integrations are common. Southern blot hybridization analyses characterized locus-specific targeting of the following mouse gene loci: *S100a8*, *Trek1*, *Inf2*, *Trpc6*, *Ccnd2* (Fig. 2E, Fig. S4-7). In all cases, ^32^P-labeled donor DNA templates were utilised as a specific-probe for hybridization (Table S2). In order to facilitate the correct detection of single copy integrations, we incorporated additional restriction endonuclease recognition sites adjacent to the introduced *LoxP* sequences (Fig. 2B, Fig. S4-7A). Restriction endonuclease recognition sites were chosen depending on the presence of the same sites in the targeted locus, assuming that after digestion of genomic DNA, the resulting fragments would be unambiguously identifiable by size during electrophoresis in 0.8% agarose gels. For example, for the desired *S100a8* conditionally targeted locus, the flanking *Bam*HI endonuclease sites were located 4 kb apart in the wild-type allele (Fig. 2A). Complete digestion of genomic DNA of the correctly targeted locus should reveal 3.2 kb, 0.7 kb and 0.3 kb DNA fragments (Fig. 2C), while the observed 1.1 kb and 0.2 kb fragments indicated multiple head-to-tail integrations of donor DNA via the NHEJ-HDR mechanisms (Fig. 2D). Using this strategy, we could clearly identify multiple copy integrations of donor DNA template during the generation of conditional KO mouse models, both in F0 or F1 offspring (Table 1, Fig. 2E, Fig. S4-7). Our analyses further indicated that multiple head-to-tail donor DNA template integrations arose via HDR-NHEJ, the HDR-HDR or NHEJ-NHEJ mechanisms (Table 1, Fig. 3, Fig. S4-7). Overall, we conclude that the repetitive head-to-tail integration of the donor DNA template is a common by-product of the CRISPR/Cas9-mediated HDR-based genome editing process, regardless of the donor DNA template size, sequence composition or strandedness of the template (dsDNA or ssDNA) (Table 1). Remarkably, Southern blot hybridization analysis enabled identification of single copy, positively targeted mice already in the F0 generation (Fig. S7, Table 1). However, due to the mosaic nature of donor DNA integration for some of the F0 mice, which indicated multiple copy integrations, we were— after crossing—able to identify offspring that harboured the desired single copy targeted allele.

## Discussion

Within a short time, CRISPR/Cas9 endonuclease has emerged as a state of the art tool for genome editing in model organisms from all kingdoms of life ^26^. From the assembly of the CRISPR/Cas9 complex and the discovery of direct targeting of specific genomic sequences *in vitro* ^9, 27^, it took only six months to experimentally verify *in vitro* findings in bacterial and mammalian cells ^3, 4, 28^. The establishment of genetically modified mouse models to study the potential functional roles of genes and their products in human diseases is an important aspect of biomedical studies ^29–33^. Conditional gene knockout mouse models constitute a powerful approach that enables the investigation of gene functions in specific cell-types and/or in a development-specific manner ^34, 35^.

Nevertheless, our study uncovered serious pitfalls exemplified in ten separate knock-in procedures during the construction of six conditional KO mouse models that need to be taken into account. All gene-targeting protocols were performed by direct injection of CRISPR/Cas9 components together with donor DNA template into fertilized oocytes. Eight KIs were performed with relatively long donor DNA fragments (∼700 – 1650 nt). Seven procedures employed ssDNA and three dsDNA templates (Table 1). Three KI attempts with ssDNA and one with dsDNA templates did not yield the desired single copy integration of donor template (Table 1).

Efficiencies of donor DNA integration were variable and correlated with template size, whereby, in general, longer templates integrated less efficiently (Table 1). We noticed that most edited mice obtained from CRISPR/Cas9 modified zygotes (F0 generation) exhibited mosaic genotypes, harbouring subpopulations of cells derived from different DNA integration events, and contained diverse copy numbers in the targeted loci. Our data suggest that PCR amplification of short genomic flanking regions together with parts of an insert is the most efficient and reliable approach for the identification of F0 mice with a correctly targeted locus. Positive PCR results on both flanks indicated that a certain subpopulation of cells contains HDR integrated DNA template (Fig. 1C, D). However, longer PCR products representing subpopulations of cells with target DNA integrated via HDR-NHEJ or NHEJ-NHEJ are difficult to amplify. Nevertheless, in some cases, most probably depending on the degree of mosaicism and PCR primer locations, such arrangements could be detected as well (Fig. 1D, numbers 10, 18).

When the selected F0 founders were crossed with wild type mice for production of the F1 offspring, we often detected animals harbouring multiple head-to-tail integrations of the donor template at the targeted loci (Fig. 3). We observed template multiplication irrespectively of the size, nucleotide composition or the utilization of dsDNA or ssDNA (Table 1). Importantly, a commonly applied PCR verification method in heterozygotic animals using template specific primers in most cases erroneously identified those as single copy integration events. Moreover, in cases of multiple-copy HDR-HDR based integrations of donor DNA, it proved impossible to correctly identify the desired single-copy mice by amplification with primers set in the genomic flanking regions followed by PCR product sequencing.

To correct this error, we propose methods that can be used for the successful identification of HDR-HDR based single copy targeted mouse loci. The first approach is based on a combination of PCR analyses: F0 and F1 founders harbouring an HDR-HDR based insertion of donor DNA could be identified using PCR amplification of flanking regions including part of the insert (Fig. 1C, D). A repeated head-to-tail template could be detected by a second PCR step using bidirectional, non-overlapping primers (Fig. 4C). Furthermore, candidates for singly targeted loci should be sequenced to confirm the absence of possible mutations in the inserted donor DNA template. This relatively simple strategy could be useful for verification of any genome knock-in models including point mutations in genes, specific deletions or insertions in all species. Notably, identification of F0 founders with positive PCR results on both flanks does not guaranty that their offspring will contain the correctly targeted single copy locus. On the other hand, identification of single copy positively targeted mice in the F0 generation is relatively rare. Since the mosaic nature of donor DNA integration often results in subpopulations of germ cells with correctly targeted loci, we therefore recommend crossing F0 candidates displaying HDR-HDR integrated donor DNA template with wildtype animals and to perform a second PCR step using bidirectional, non-overlapping primers on F1 offspring.

As shown in this study, Southern blot analysis is an additional method to reliably identify the desired F1 founders. Below, we outline a strategy to design donor templates that permits the unambiguous identification of single-copy targeted loci. We recommend the incorporation of two specific restriction endonuclease sites flanking the *LoxP* regions. This will allow the detection of small DNA fragments on Southern blots in the event that multiple donor template copies are integrated (Fig. 2E, Fig. S4-7). Notably, the fragments should not be too small, as Southern blots are unable to detect small numbers of repeats.; this is exemplified by the failure to expose the 0.2 kb signal in the *Trpc6* gene conditional KO project (Fig. S6C).

Despite the advantages of CRISPR/Cas9 based genome editing, a number of potential problems such as target specificity and off target effects still impede the CRISPR/Cas9 technology for use in biomedical research and further efforts are necessary to overcome these hurdles. Our study examines problems that are not unique for the CRISPR/Cas9 system, but instead generally affect direct knock-in genome targeting. In multiple cases, we documented that the insertion of donor DNA via the HDR mechanism results in mosaicism yielding sub-populations of cells with head-to-tail template amplification in the modified loci. Our findings are important to unlock the full potential of the CRISPR/Cas9-mediated genome editing protocols for the generation of custom designed gene variants for biomedical research and gene therapy.

## Materials and Methods

### Cytoplasmic microinjections of the CRISPR/cas9 components into fertilized oocytes

For the preparation of CRISPR/Cas9 microinjection solution, commercially synthesized crRNA (Table S1), tracrRNA and Cas9 protein (IDT, USA) were mixed as follows: 100 pmoles of crRNA were mixed with 100 pmoles of tracrRNA (when two crRNAs were used, the concentration of tracrRNA was increased to 200 pmoles) in 10 mM potassium acetate, 3mM Hepes (pH 7.5) buffer and incubated at 95 ^0^C for 2 minutes following by cooling to room temperature. The annealed crRNA/tracrRNA complex was mixed with Cas9 mRNA, Cas9 protein and DNA injection fragment. The final concentrations of CRISPR/Cas9 components in 0,6 mM Hepes (pH=7.5), 2 mM potassium acetate microinjection buffer were: 2 pmol/µl of crRNA, 2 pmol/µl of tracrRNA (or 4 pmol/µl of tracrRNA if two crRNAs were used), 10 ng/µl of Cas9 mRNA, 25 ng/µl of Cas9 protein and from 0,05 to 0,01 pmol/µl DNA target fragment. The final injection solution was filtered through Millipore centrifugal columns and spun at 20,000g for 10 min at room temperature.

Microinjections were performed in B6D2F1 (hybrid between C57Bl6/J and DBA strains) fertilized one-cell oocytes. Oocytes were removed from oviducts of superovulated B6D2F1 female mice in M2 media supplemented with hyaluronidase (400 μg/ml), washed twice for removal of cumulus cells in M2 media, transferred to KSOM media, and kept at 5% CO2 and 37°C before injections. Cytoplasmic microinjections were performed in M2 media, using the Transjector 5246 (Eppendorf), and Narishige NT-88NE micromanipulators attached to a Nikon Diaphot 300 inverted microscope. Oocytes that survived microinjections were transferred to oviducts of pseudopregnant CD1 foster mice and carried to term. Positively targeted F0 animals were identified by PCR and Southern blot analysis of genomic DNA isolated from tail biopsies.

### Donor DNA template preparation

Donor DNA templates for microinjection (Table S3) were synthesized and cloned into pUC57 or pBlueScript vector (Biomatic). dsDNA templates were sequenced and directly digested from the CsCl_2_ gradient purified plasmid vector using *Xho*I restriction endonuclease. The resulting donor dsDNA fragments were separated using 1% agarose gel electrophoresis, extracted with 6M NaI and stored in ddH_2_O. ssDNA templates were either purchased from IDT or MWG or amplified from the aforementioned plasmid vectors using asymmetric PCR with 500 molar excess of one of the primers. PCR amplification was performed in 50 µl reaction volume containing 200 ng of plasmid DNA template, 1 pmol/µl and 0,002 pmol/µl of primers (Table S3), 50 U of Taq polymerase, 2 U of Phusion DNA polymerase (NEB) and 0,2 mM dNTPs. The resulting ssDNA fragments were separated using 1% agarose gel electrophoresis, extracted with 6 M NaI and stored in ddH_2_O.

### PCR analysis of the targeting events for HDR, NHEJ and multiple copy integration

PCR analysis was performed in 50 µl reaction volume containing 1 µM of each gene specific primer (Table S3), 5 U of Taq polymerase, 100 ng of genomic DNA, 5% DMSO, 1 M Betain and 0,2 mM dNTPs. The resulting DNA amplicons were separated using 1% agarose (1X TAE buffer) or 6% (w/v) polyacrylamide gel (1 X TBE buffer) electrophoresis, followed by ethidium bromide staining.

### Southern blot DNA analysis

Genomic DNA was obtained from tail biopsies. Tail tissue was lysed in buffer containing 100 mM Tris-HCl, pH 8.5; 5 mM EDTA; 0.2% sodium dodecyl sulfate (SDS); 200 mM NaCl; and 100 µg/ml proteinase K (Roche) overnight at 55°C. Genomic DNA was extracted by phenol-chloroform and chloroform followed by precipitation with 2.5 volumes of isopropanol and washing with 70% ethanol. The DNA pellet was dissolved in TE buffer (10 mM Tris, pH 7.9; and 0.2 mM EDTA). Positively targeted F1 animals were analyzed using Southern blot hybridization. Approximately 10 - 20 µg of genomic DNA was digested with the corresponding restriction endonuclease, fractionated on 0.8% agarose gels, and transferred to GeneScreen nylon membranes (NEN DuPont). The membranes were hybridized with ^32^P-labeled specific DNA probes (Table S2). DNA labelling was performed using random prime DNA labeling kit (Roche), and [α-^32^P] dCTP (PerkinElmer). Membranes were washed with 0.5x SSPE (1x SSPE is 0.18 M NaCl, 10 mM NaH_2_PO_4_, and 1 mM EDTA, pH 7.7) and 0.5% SDS at 65°C and exposed to MS-film (Kodak) at −80°C.

### Mice

All animal procedures were performed in compliance with the guidelines for the welfare of experimental animals issued by the Federal Government of Germany. F1 heterozygous mice were produced by breeding F0 DBAxC57Bl/6J founders to C57Bl/6J mice.

Pups were weaned at 19 to 23 days after birth, and females were kept separately from males. The mice were housed in standard individually ventilated cages (IVC). General health checks were performed regularly in order to ensure that any findings were not the result of deteriorating physical conditions of the animals.

## Acknowledgments

We thank Stephanie Klco-Brosius for help with editing. This work was supported by the Deutsche Forschungsgemeinschaft (RO5622/1-1 to TSR, SK259/2-1 to BVS, SU 195/4-1 to CS, SFB/CRC 1009-B09 to JR and SFB/CRC 1009-B10 to HP and RWS), BMBF (16GW0055 to SGM), DZHK (81X2800174 to BVS), the IZKF Muenster (RWS, Wed2/022/18), Cancerfonden (CAN 2016/524 to YBS), Strategic Research grant from Medical Faculty, Umeå University (to YBS) and an IMF grant of the Medical Faculty of WWU (SH121608 to JS). Core Facility TRAM is an institution of the Medical Faculty of WWU. The support of the Medical Faculty is thankfully recognized.

## Competing interests

The authors declare no competing interests.

## Author contributions

BVS and TSR conceived and designed the study. LG, BS, HK, AS, BVS and TSR were involved in experimental work for all cKO projects. JR, SGM, YBS and CS were involved in generation of *S100a8*, *Trek1*, *Ccnd2* and *Il4* cKO mice, respectively. HP and RWS participated in *Inf2* cKO project. JS and TP were involved in *Trpc6* cKO mice generation. TSR, BVS and JB analysed data and wrote the paper. All authors provided input and approved the final manuscript.

