## Supplementary Material and Methods; Tables S1-3; Figures S1-7 for "Pervasive head-to-tail insertions of DNA templates mask desired CRISPR/Cas9-mediated genome editing events"

[1]Medical Faculty, Core Facility Transgenic animal and genetic engineering Models (TRAM), University of Muenster, Muenster, Germany; [2]Institute of Immunology, University Hospital Muenster, Muenster, Germany; [3]Clinic of Neurology with Institute of Translational Neurology, University Hospital Muenster, Muenster, Germany; [4]Internal Medicine D, University Hospital Muenster, Muenster, Germany [5]Institute of Experimental Musculoskeletal Medicine (IMM), University Hospital Muenster, Muenster, Germany; [6]Institute of Cell Dynamics and Imaging, University of Muenster, Muenster, Germany; [7]Department of Dermatology and Venereology, University Hospital Halle, Martin Luther University Halle-Wittenberg, Halle (Saale), Germany; [8]Department of Molecular Biology, Umeå University, 901 87 Umeå, Sweden, [9]Institute of Experimental Pathology (ZMBE), University of Muenster, Muenster, Germany; [10]Institutes for Systems Genetics, West China Hospital, Sichuan University, Chengdu 610041, China; [11]Brandenburg Medical School (MHB), Neuruppin, Germany.

### Supplementary Material and Methods

#### *Quantitative PCR (qPCR) analysis*

TaqMen qPCR analysis was performed using 100 ng of genomic DNA samples. All details on the specific primer sequences and dual labeled LNA probes (IDT)) used for qPCR analysis are provided in Table S3. All qPCR reactions were performed in duplicates in a total volume of 20 µl containing 2 µl of genomic DNA (~100 ng), 10 µl of 2 X TaqMan Master Mix (Roche), 0,1 µM TaqMan LNA probe and 0,1 µM of each primer. PCR amplification was performed as follows: enzyme activation at 95C for 5min (Ramp Rate (RR) 4,4), with subsequent 50 cycles of qPCR: 95C – 10sec (RR 4,4); 60C – 30sec (RR 2,2) and final cooling cycle: 40C – 10sec (RR 2,2). Quantification Cycle (Cq) values were calculated using Light-cycler 480 SW 1.5 software E-method (Roche) and all data were further analyzed with Excel.

#### Analysis of DNA template copy number integration using digital droplet PCR (ddPCR)

Genomic DNA from F1 mice (derived from *S100A8* founder Nr.6 and Nr.11) was diluted up to 10 ng/µl for all ddPCR experiments. The primers pair dd5LoxP_Dir and dd5LoxP_Rev (Table S3) (final concentration, 100 nM) and 5'-6-FAM/3'IBFQ-labeled probe (AACCTACTTGAGGGCCCACT) were used for the *S100A8* 5'-*LoxP* assay. As a reference dHsaCP1000001 HEX-labeled assay (Bio-Rad Laboratories) was utilized. ddPCR reactions were performed in a total volume of 20 µl containing 20 ng of genomic DNA and ddPCR EvaGreen Supermix (Bio-Rad Laboratories, Hercules, CA, USA). All reactions were placed in eight-well disposable cartridge (DG8™; Bio-Rad Laboratories) together with 70 µl of droplet generation oil (Bio-Rad Laboratories) for production of droplets using a QX200™ Droplet Generator (Bio-Rad Laboratories). Subsequently, samples were PCR amplified using Biometra TOne thermal cycler (Analytik-Jena) as follows: enzyme activation at 95 °C for 5 min, followed by 40 cycles of 94 °C for 30 s and 60 °C for 1 min, and final signal stabilization at 98 °C for 10 min). The resulting droplet PCR products were analyzed using the QX200 Droplet Reader (Bio-Rad Laboratories), and QuantaSoft™ software (Bio-Rad Laboratories).

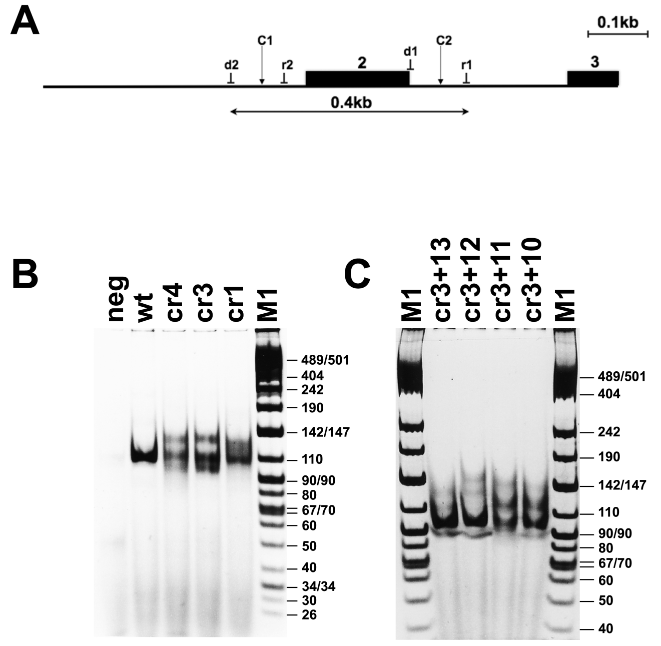


**Supplementary Figure 1**. Evaluation of *in vivo S100A8* crRNA cleaving efficiency in mouse embryos. (**A**) Schematic representation *S100A8* targeting region. The intronic regions are shown as line, and exons are drawn as filled boxes numbered above. The arrows above the line correspond to target site regions for the CRISPR/Cas9 complexes with either crRNA-10, crRNA-11, crRNA-12, or crRNA-13 (C1), and crRNA-1, crRNA-3, or crRNA-4 (C2), respectively. The positions of direct (d1, d2) and reverse (r1, r2) orientation PCR primers are denoted. (**B, C**) PCR analysis of DNA samples isolated from 30-60 two-cell embryos on the day after microinjections of the CRISPR/Cas9 complexes using 6% (w/v) polyacrylamide gel (1 X TBE buffer) electrophoresis, followed by ethidium bromide staining. PCR amplification was performed with (**B**) d1-r1 and (**C**) d2-r1 primer pairs. (**B**) CRISPR/Cas9 RNP was assembled with either crRNA-1, crRNA-3, or crRNA-4 (cr1, cr3, cr4). The DNA size standards (M1) are presented on the right in bp. The size of the DNA fragment corresponding to the PCR product of the unmodified locus is 105 bp (wt). Samples cr3 and cr4 feature additional, lower size PCR products, corresponding CRISPR/Cas9 RNP cleavage; "neg."denotes negative control (PCR without DNA). (**C**) CRISPR/Cas9 complexes either contained crRNA-3 + crRNA-10, crRNA3 + crRNA-11, crRNA-3 + crRNA-12, or crRNA-3 + crRNA-13 (cr3+10, cr3+11, cr3+12, cr3+13) crRNA pairs. All samples exhibit a PCR product about 100 bp in size, which corresponds to the predicted deletion fragment.

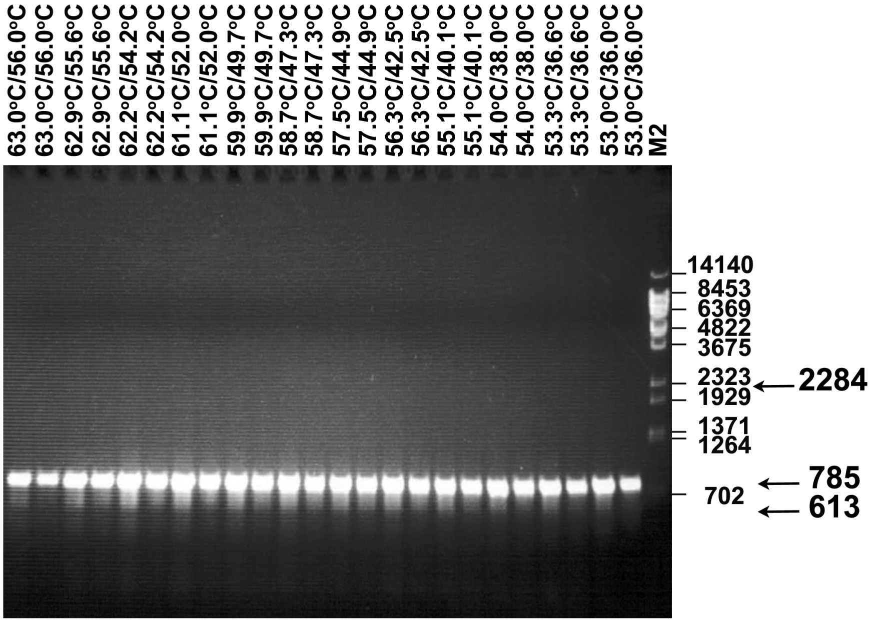


**Supplementary Figure 2**. PCR analysis of genomic DNA from F0 founder number 6 after HTTP integration in the *S100A8* locus at different touch down/annealing temperature conditions using primer pair (d4/r4) (Figure 1D). The PCR product (785 bp) corresponds to the single copy correctly targeted *S100A8* gene. The PCR product corresponding to the wt allele (613 bp) is not detected. PCR products (2284 bp) indicating multiple head to tail integrations of the DNA template is also undetectable.

**A**

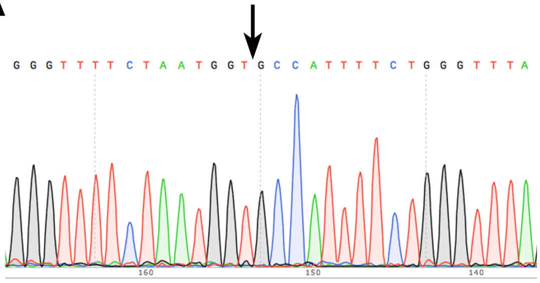


**B**

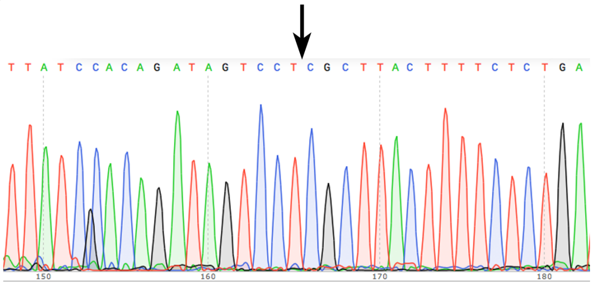


**Supplementary Figure 3.** Sequence analysis of heterozygous animal (F1) number 45 with MC head to tail integration of the DNA template in the *S100A8* gene (Figure 1E, 2A). (**A**) Sequence analysis of the left flanking region. The MC DNA template integrated via NHEJ after cleavage of the *S100A8* gene locus at the *C2* site (Figure 1E). The arrow indicates the fusion point. (**B**) Sequence analysis of the right flanking region. The MC DNA template integrated via HDR after cleavage of the *S100A8* gene locus at the *C2* site (Figure 1E). The arrow indicates the junction between the DNA template homologous and non-homologous sequence.

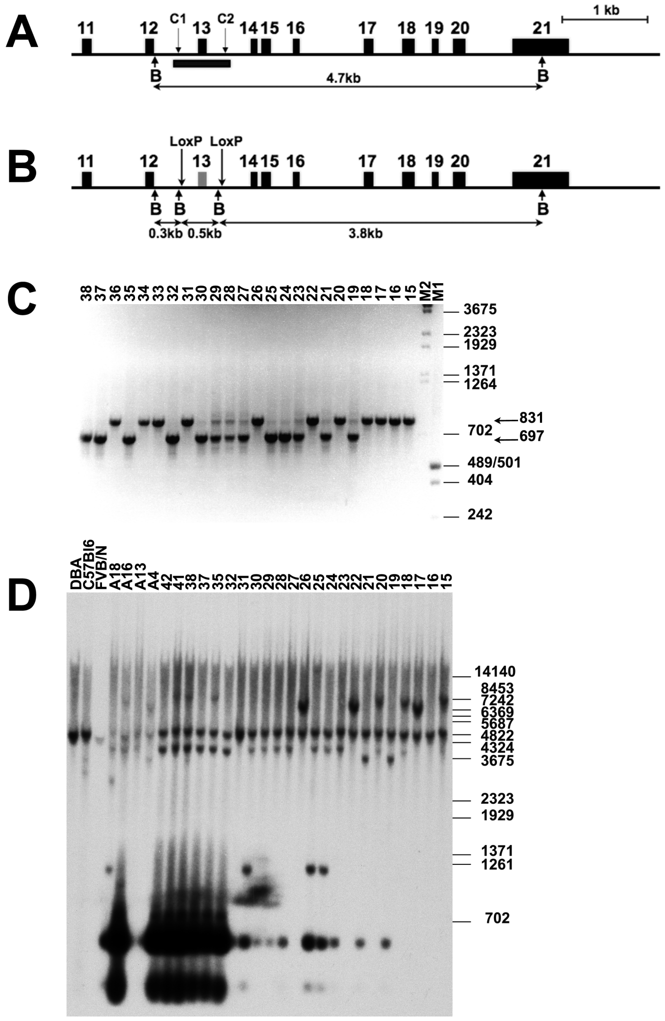


**Supplementary Figure 4**. Analysis of the *Inf2* targeted locus. (**A**) Schematic representation of the *Inf2* gene targeting strategy. Exon 13 was selected for elimination. Intronic and intergenic regions are shown as line, exons are shown as filled boxes numbered above. The arrows above the line correspond to the target sites of the CRISPR/Cas9 complex with crRNA 15 (C1) and crRNA 7 (C2), respectively. The arrows marked "B" correspond to *Bam*HI restriction endonuclease sites. The black bar below exon 13 corresponds to areas recognized by the donor DNA specific probe used in Southern blot analyses. The horizontal arrows denote the expected sizes of restriction DNA fragments given in kb. (**B**) Schematic representation of the *Inf2* gene targeted locus. Positions of inserted *LoxP* sites are indicated by vertical arrows. (**C**) PCR analysis of genomic tail biopsy DNA samples from F1 mice 15-38 (labeled above), using primer pair (d2/r1) located at the ends of the DNA template homology arms (not shown). The PCR products for correctly targeted (697 bp) and wild type (831 bp) alleles are labeled by arrows. PCR products corresponding to multiple head-to-tail integrations of the DNA template were not detected. Size marker positions (in bp) are shown on the right. (**D**) Southern blot analysis of genomic DNA samples hybridized with the template specific probe denoted in (**A**) by the black bar. DNA samples were obtained from mouse-tail biopsies of F1 offspring: 15-22, 26 {F0 A4}; 23-25, 27-30 {F0 A12}; 31, 32, 35, 37, 38, 41, 42, {F0 A16}; and F0 mice: A4, A13, A16, A18. *FVB/N*, *C57Bl6* and *DBA* are wt DNA samples from the corresponding mouse lines. Notably, the same F1 offspring DNA samples were used in PCR analysis. *Bam*HI enzymatic digestion detected the wild-type allele (4.7 kb) and three DNA fragments (3.8, 0.5 and 0.3 kb) corresponding to the targeted allele (B). DNA samples 28, and 29 contain the correctly targeted *Inf2* allele (*Inf2*$^{+/-}$). Samples 24, 25, 31, and A18 contain DNA fragments of 1.1kb size, indicating insertion of the MC head to tail integration through the NHEJ at the left flank and HDR at the right flank at the C2 site (Figure 3B4). Samples 32-42 show MC head to tail integration of the DNA template through the HDR-HDR mechanism (Figure 3B2). Size marker positions (in bp) are shown on the right.

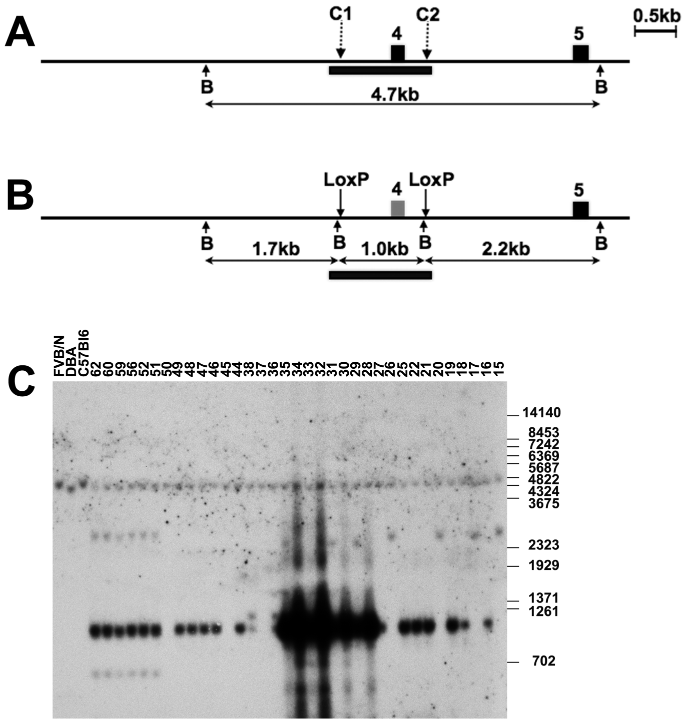


**Supplementary Figure 5**. Analysis of the *Trek1* targeted locus. (**A**) Schematic representation of the *Trek1* gene targeting strategy. Exon 4 was selected for elimination. Intronic and intergenic regions are shown as line, exons are shown as filled boxes numbered above. The arrows above the line correspond to the target sites of the CRISPR/Cas9 complex with crRNA-2 (C1) and crRNA-4 (C2), respectively. The arrows marked "B" correspond to *Bam*HI restriction endonuclease sites. The black bar below exon 4 indicate a region recognized by the donor DNA specific probe used in Southern blot analyses. The horizontal arrows denote the expected sizes of *Bam*HI restriction DNA fragments given in kb. (**B**) Schematic representation of the *Trek1* gene targeted locus. Positions of inserted *LoxP* sites are indicated by vertical arrows. (**C**) Southern blot analysis of genomic DNA samples from mouse-tail biopsies of F1 mice: 15-22, 25-38, 44-52, 56, 59, 60, 62 and *FVB/N*, *C57Bl6* and *DBA* lines of wild type animals, hybridized with the template specific probe (A). *Bam*HI enzymatic digestion detected the wild-type allele (4.7 kb) and a DNA fragment (1.0 kb) corresponding to the targeted allele (B). DNA samples 16, 18, 19, 21, 22, 25, 27-35, 44, 46-49, 51, 52, 56, 59, 60 and 62 contain the multiple head to tail integrations of the DNA template (MC) at the targeted locus. Samples 51, 52, 56, 59, 60 and 62 (F0 founder 8) reveal a DNA fragment of 2.7kb, indicating insertion of the MC head to tail via NHEJ at the left flank and HDR at the right flank at the C2 site (Figure 3B4). Samples 44, 46-49 (F0 founder 8) confirmed by sequencing MC integration through the HDR mechanism at both sites C1 and C2 (not shown) (Figure 3B2). Size marker positions (in bp) are shown on the right.

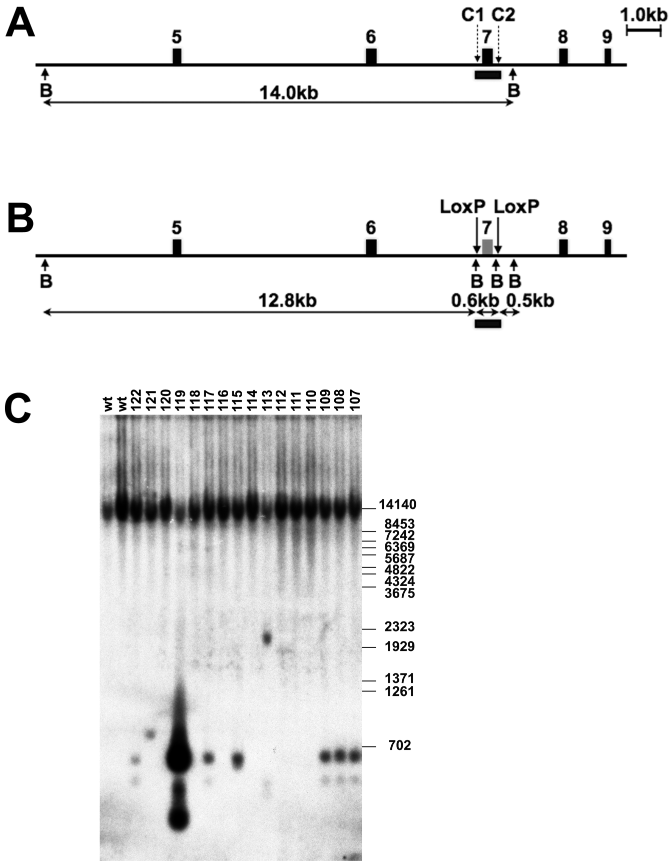


**Supplementary Figure 6**. Analysis of the *Trpc6* targeted locus. (**A**) Schematic representation of the *Trpc6* gene targeting strategy. Exon 7 was selected for elimination. Intronic and intergenic regions are shown as line, exons are shown as filled boxes numbered above. The arrows above the line correspond to the target sites of the CRISPR/Cas9 complex with crRNA-1 (C1) and crRNA-5 (C2), respectively. The arrows marked "B" correspond to *Bam*HI restriction endonuclease sites. The black bar below exon 7 depicts a region recognized by the donor DNA specific probe used in Southern blot analyses. The horizontal arrows denote the expected sizes of restriction DNA fragments given in kb. (**B**) Schematic representation of the *Trpc6* gene targeted locus. Positions of inserted *LoxP* sites are indicated by vertical arrows. (**C**) Southern blot analysis of genomic DNA samples from mouse-tail biopsies of F1 mice: 107-122, and wild type (wt) animals, hybridized with the template specific probe (A). *Bam*HI enzymatic digestion detected the wild-type allele (14 kb) and two DNA fragments (0.6 and 0.5 kb) corresponding to the targeted allele (B). DNA samples 107-109 contain the correctly targeted *Trpc6* allele (*Trpc6* $^{+/-}$). Sample 119 contains strong signals corresponding to 0.6 kb and 0.2 kb DNA fragments, indicating MC head to tail integration via HDR mechanism at the C1 and C2 sites (Figure 3B2). Sample 115 contains a slightly enhanced signal for the 0.6 kb DNA fragment but is devoid of 0.2 kb signal presumably due to the small size of the fragment and the low degree of MC. Size marker positions (in bp) are shown on the right.

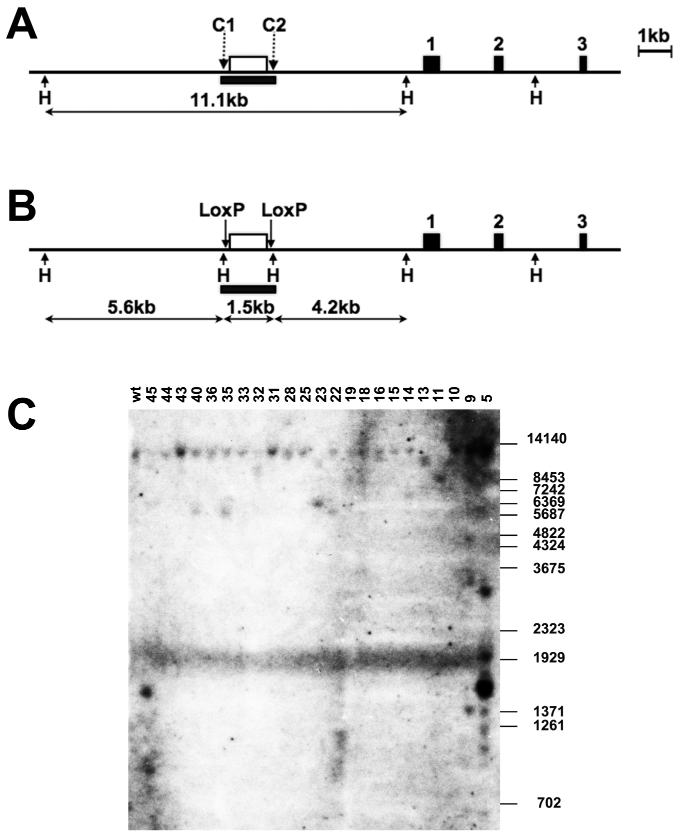


**Supplementary Figure 7**. Analysis of the *Ccnd2* targeted locus. (**A**) Schematic representation of the *Ccnd2* gene targeting strategy. The PTE regulatory region (open box above the line) was selected for elimination. Intronic and intergenic regions are shown as a line, exons are shown as filled boxes numbered above. The arrows above the line correspond to the target sites of the CRISPR/Cas9 complex with crRNA-1_2 (C1) and crRNA-2_2 (C2), respectively. The arrows marked "H" correspond to *Hind*III restriction endonuclease sites. The black box below the targeted are corresponds to a region recognized by donor DNA specific probe used in Southern blot analyses. The horizontal arrows denote the expected sizes of *Hind*III restriction DNA fragments given in kb. (**B**) Schematic representation of the *Ccnd2* gene targeted locus. Positions of inserted *LoxP* sites are designated by vertical arrows. (**C**) Southern blot analysis of genomic DNA samples from mouse-tail biopsies of F0 mice: 5, 9, 10, 11, 13-16, 18, 19, 22, 23, 25, 28, 31, 32, 33, 35, 36, 40, 43-45, and wt animals, hybridized with the template specific probe (A). Southern blot analysis detected the wild-type (11.1 kb) and correctly targeted (1.5 kb) alleles in DNA sample 45 (*Ccnd2* $^{+/-}$) (verified by sequencing of the F1 offspring (not shown)). DNA sample 5 contains the multiple head to tail integration of the DNA template (MC) at the targeted locus. Size marker positions (in bp) are shown on the right.

**Supplementary Table 1.** List of crRNAs used.

| Gene (template) name | crRNAs |
|---|---|
| *S100A8* | crRNA3 GTTTTCTAATGGTGCTGGGA<br>crRNA12 TCAGGTTTTTAACCCAAGAA |
| *Trek1* | crRNA2 GACATCCCGAACAGGTGTAG<br>crRNA4 GGGCTCCACTTGAAGTTGTA |
| *Inf2* | crRNA7 GGTCCTGAGCACTTGGGGT<br>crRNA15 GGGCTGCTACCAAGTTGCC |
| *Trpc6* | crRNA1 GAGAGAGAAATGAACCTGAG<br>crRNA5 TTGCAGGAAGACTAGCATAA |
| *Ccnd2* | crRNA1_2 ACTGTGCTCTGGAACCATCC<br>crRNA2_2 CACAAGTACGGATGCCACGT |
| *IL4_5'LoxP* | crRNA10 GGAATGAGGCTTTACCTATA |
| *IL4_flox* | crRNA1 GACTCCCTCTGCCTCCAAGG<br>crRNA10 GGAATGAGGCTTTACCTATA |

**Supplementary Table 2.** Designed donor DNA templates. Notably, same DNA templates were used as gene specific-probes in Southern blot hybridization.

| **Gene (template) name** | **Nucleotide sequence (5’-3’)** |
|---|---|
| *S100A8* | AGCCATTTTCTGGGTTTAGTTATCTGAGTGGTACAGGCCAGGTCAAAGCTTATTCTTGGGTT<br>AAAAACCTGAATGCGGATCCAAGCTGGGCCCTGGTAATATAACTTCGTATAGCATACATTA<br>TACGAAGTTATGGCTGCGGAAACAACCTACTTGAGGGCCCACTCAGTTGTCACCCTGCCGT<br>GAGTAACTGCAGCTCCCATCCCTCTGCGACTTTTCCTTTCAGTTGAAAGGAAATCTTTCGTG<br>ACAATGCCGTCTGAACTGGAGAAGGCCTTGAGCAACCTCATTGATGTCTACCACAATTATT<br>CCAATATACAAGGAAATCACCATGCCCTCTACAAGAATGACTTCAAGAAAATGGTCACTA<br>CTGAGTGTCCTCAGTTTGTGCAGGTGAGGAGGTGCTATGTTCGGATCCAAGCTGGGCCCTG<br>GTAATATAACTTCGTATAGCATACATTATACGAAGTTATGGAGAGGAGGAAACAACCTAC<br>TTGAGGGCTCAGGGTTTTCTAATGGTGCTGGGATTTCTAGCCCTCCAAGTTGCTTTCTGCAG<br>TTAGCCAATACTTATCCACAGATAGTCCT |
| *Trek1* | AGAGACAAAGGATTATCTACCATACCTGTGTTCTCTGTGCGTCTGGGACATCCCGAACAGG<br>TGTAGGTGGGATCCAAGCTGGGCCCTGGTAATATAACTTCGTATAGCATACATTATACGAA<br>GTTATGGCTGCGGAAACAACCTACTTGAGGGCATACCTCAGCAATTATGAGGTCCGTGAA<br>AGGCAATGGCCTCCATCATGGCAAGTTCATGTCTTTGGACCTTGAAACTCTTCAGAATGTC<br>ACATTTTGGCATTTATCTATCTTTAAAATATAATAGGATTTTAAAAAAAGACATAAATTGT<br>AACACTGCCTCACTCTTTAGATAATGTACGTGTGTTTGAAGATTCTTAATGTATCTCTCTGA<br>ACACATTATAAATGGTAATGTAAGTTTTGTAAGTCAATAGCAATTATCATTGCCTGGGAGA<br>CAGAAGGAGACTAATGAAAATATTGACTCCATTAGGTGCGTTAACACCGCATTGCTCATCG<br>TCTGATCAAAGTTTCACCAAACATTAGCTGGATGTGGTATTTTATTAGAATAATTGCTTAC<br>ATTAGCTTTTACTTGTACTGTTATAATATAAATTGAGTCTTCCCCAGGTGAAATAAACAATC<br>CAGGTCTTGTAATAGCTGGTGATTTCTGGTTAAACTTGGCATGGGGTGTTTAAAACTATGA<br>TACATTTGTAAATAACTGGATTTTTCCATTTTTGGTTTTCTTTTTAGGATTTGGAAACATCTC<br>CCCACGAACTGAAGGTGGAAAAATATTCTGCATCATCTATGCCTTGCTGGGAATTCCCCTC<br>TTTGGCTTTCTACTGGCTGGGGTTGGTGATCAGCTAGGAACTATATTTGGAAAAGGAATTG<br>CCAAAGTGGAAGACACATTTATTGTGAGTAGCACAAACTTCTTGCTACATCTATTTAATGG<br>TTTTGAAAATATGTTACATATTCTAGCACCTTAGATAATGTAGACCTTCAGTTGCCTTTTAT<br>ATAACAATTTCTATGCCTCTTAGCTTGCCACTAACTGTGTTTGACTTATTTACATACTTGAT<br>TGAAAGAAACTGAAAATTATGTACAAAAAAACAGAATTATAACCTTGGATCCAAGCTGGG<br>CCCTGGTAATATAACTTCGTATAGCATACATTATACGAAGTTATGGAGAGGAGGAAACAA<br>CCTACTTGAGGGCCCTGGGCTCCACTTGAAGTTGTATTTTGTCCTAATTATTTTCTCTGTGT<br>GTGTGTGCACATGCAAACTTG |
| *Inf2* | CCCTGAGGCCAGAGACTTCCTCAGAAGCTCCAGGCATCCTGGGCTGCTACCAAGTTGCCCT<br>TGCCTGACTTCGGATCCAAGCTGGGCCCTGGTAATATAACTTCGTATAGCATACATTATAC<br>GAAGTTATGGCTGCGGAAACAACCTACTTGAGGGCTGTCCTGGGCACAGGCCTAGGAATG<br>TATATGTTCTCAGCCCAGGCTATGGGGACTCTCCCTCAGCCCACCCACCAAGGGCAGAGCA<br>TGTTCCCAGTGGCAAGCATGGTGGAACCTCCAGGCCCTAACCCCCTCCTGTGCACCAGCTA<br>CCCGCTGCGGGTGGAGTGCATGATGCTGTGTGAGGGAACGGCCATCGTCCTGGACATGGT<br>GCGGCCCAAGGCCCAGCTGGTGCTCACTGCCTGCGAGAGTGAGTGGGGCCCAGAAGCCTA<br>CGGGGCTGGGAGGGATGGGCCTTCCACGTGTGTCAGCCCCTGCTGGAGCCATGTCACAAT<br>GGCCACGTGAGGTGCCAGCTTTGCCACCGAAACCCATAGTCACCTCTGCTGCTAACACAGG<br>ATCCAAGCTGGGCCCTGGTAATATAACTTCGTATAGCATACATTATACGAAGTTATGGAGA<br>GGAGGAAACAACCTACTTGAGGGCGGGTCCTGAGCACTTGGGGTGTTGGGGTGCCACAGA<br>CTGGAAGGAGTTCCTGTAGCATCCCAGAGTC |
| *Trpc6* | GAGAGCCACAGCAGCAGAATCACCTAATCATTGGAGAGAGAGAAATGAACCTGAGTTGTA<br>ATTTGCATATATGGATCCAAGCTGGGCCCTGGTAATATAACTTCGTATAGCATACATTATA<br>CGAAGTTATGGCTGCGGAAACAACCTACTTGAGGGCCACACTTGAGAAGTTCTTCAGAGC<br>ACAAGGTTTTACTTGGGAATTGTGATTATACATTTATTTTCTCAGAAAAGGATTTACCTAA<br>ATAAACAACTCATGGGAACTTACCATAATCTTTGTCTCTGATTTTTTATCTCCAGCCAGGAT<br>AAAGTGGGACCCTACTGATCCTCAGATCATCTCTGAAGGTCTTTATGCAATCGCTGTGGTT<br>TTAAGTTTCTCCAGAATAGCTTACATTTTACCAGCAAATGAAAGCTTTGGACCTCTGCAGA<br>TTTCACTTGGAAGAACAGTGAAAGATATCTTCAAATTCATGGTCATATTCATCATGGTGTT<br>TGTAGCCTTTATGATTGGAATGTTCAACCTTTACTCCTACTACATTGGCGCAAAACAGAAT<br>GAAGCATTCACAACGTATGTGGTGCTCGGGGTCCTGATGGTAGAGAATGCAGATAGGAGG<br>GCATTTGACCACTAGCATTTATGCTTGCAGGTTTATACTTTGATGCAGTACTCATCTCTTGC<br>TGTGTGTCAACAAGACAAGGAGATCTTAAAGTGGATCCAAGCTGGGCCCTGGTAATATAA<br>CTTCGTATAGCATACATTATACGAAGTTATGGAGAGGAGGAAACAACCTACTTGAGGGCT<br>TGCAGGAAGACTAGCATAAATCAACCTTCCCAATATAAAATAATTATATGTAATGATTTTA<br>GAGTCAGATTACATGACA |

| | |
|---|---|
| *Ccnd2* | CATACATTCCTCATAGGGGTCCGAGCTAGTGTGGGCTTAGCTTAGCATACAGTCACAAACA<br>AGCTTAAGCTGGGCCCTGGTAATATAACTTCGTATAGCATACATTATACGAAGTTATGGCT<br>GCTGCGGAAACAACCTACTTGAGGGCGAGCACAGTTTCTTCTCCAAAGGTGTACATTTGA<br>AGAATGTAAGACTTTGAAAACACCCAGCTTCAAAATGCAAAACTCAACCTTTCTTGTTTTT<br>AAATCCTGGTTAAAAAAAAAAAAAAAACAATACCAAGAAAATAATTTTAGAAGTAAATTTC<br>AACCTGCTTCTAGAAAAAAGGGTTTGGAAAGGCTAAAAGTTCAAGAAGAAAATCATTTTT<br>CCTTTAATACCACTGAATTAATTGCAGAAGGGCTGGAGAGTGAATCTGGTATACTGTGGGA<br>TTGTGGGTAAACTGTGCTAACAAATGCTGACAGTTCTTATGCTTAATGATTGAAAAATATG<br>ATAGAAAATAGAAAATACTGATGAGAATGTTTTATACGGCTGCAATTTGGTTGTAGCTGAT<br>ATATTGATACATAGCTCAATAATTACAATAAAATAGAAATGTTCAGTCACAGAGAAAGTG<br>GATGTTTTTTCAGAGGGGAAAATGTCTGCATTTGAACAAAAGTAAGTATTTAGAAAACAA<br>ATAAAATTTAACTTTTGCTTTTTAAAAAATCTGGTACGGGTAGTGTTAAATGTTTTACAGA<br>GAGAATGTCCACAATTTAAAGTGTCAAATGAAGTTGAAGCAAAATTTAAAGGAGACCTGT<br>CTTTTAGGTTCTTAAACACTTCCTTCAGTACTTTTCTTTTAAAAATGTTTAATACTGTATTCT<br>GTTTTTCTTCTTTGTATACACAAAATAAATGTTTATCACATTGGTCACCTGAAATGAGCCAT<br>TCTTCAATAAAGGTCTCACACACTGAGCCTCAAAAGCAATTTATTGACATTTTTATTAATTA<br>TAATTCCACCCCTGTCCCAGAAGTCACAAAACAAAGCTGCTTGGATCAGACTAGCAATCAT<br>TAAACGAAAAAGAACTAAATAATAAGTTAAAAATAGAGAATGTGTTCAAAGGAAGCAAG<br>AAAGTGTTGCTCATAATCTCTCTGTCCTCTTCAAAATAGACACCTTTCCAAGCCTCACCTCC<br>CATTCAAACTGAAAACTCCACTACTTTTGAAACAAATAATTCCACATCAAGTCCATTTCAA<br>AAGAAGCCCCTGGCTCCTCCCACAAGCCCCTTTACCTCGCTGAAGTTAAGAAAAAGTCAG<br>AGCCCGGTGGCATTTTTCTCCCCAGCACCCACTCTGCAGCTCCTTCAATGTGAGCATCTACT<br>CGTCTTAATTACCGTGGGCTTAAAGGAAACTGACACATTTTAGCCCCCATGCCTCCAAACT<br>GAAAGATTGAAAAGATTTTGTCCAATTTCAGGGGAAACGGAGGTGCGTGGGCTCACCCTG<br>TAGTTTTCAACCCCTGCAGTCTCTCTAGAAGGTTCTCCCAGGTTAGGATCGCATAGGATAA<br>CTTCGTATAGCATACATTATACGAAGTTATGGCTGCTGCTCCTTAATGCGCGTAGTCGAAG<br>CTTGTATCCAGTGAGTGGGCGGGGAGTGGCCCTGCACTTTCGCTCATCTCTGGTCTTCTCAC<br>A |
| *IL4_5'LoxP* | CAGCCATTTCTCAGGCTTCTGTCTAAGGTAGGAAAAATCTTCAACCTAGCCCAGAACCTCG<br>GATCCAAGCTGGGCCCTGGTAATATAACTTCGTATAGCATACATTATACGAAGTTATGGCT<br>GCGGAAACAACCTACTTGAGGGCTCGAGGTAAAGCCTCATTCCATGGTCCTGCCTGCCCCA<br>CTCCATGTCACCTCTCTGTCTCCAAAG |
| *IL4_flox* | AGCCATTTCTCAGGCTTCTGTCTAAGGTAGGAAAAATCTTCAACCTAGCCCAGAACCTTTA<br>TATAGGTAAAGCCTCATTCCGGATCCAAGCTGGGCCCTGGTAATATAACTTCGTATAGCAT<br>ACATTATACGAAGTTATGGCTGCGGAAACAACCTACTTGAGGGCATGGTCCTGCCTGCCCC<br>ACTCCATGTCACCTCTCTGTCTCCAAAGACCACAAACTTGTAAGATCAGCTGGTCTAGGAT<br>GCGAGAAGGTCTGCCTCCATCATCCTTCTATGAGGTAAGACCCCAGAGTCAGCTTTCCCAA<br>GATATCAGAGTTTCCAAGGGGCCCCCATAGCAGGAAGCAGCTAGGCCCAGGTGTGCTCAA<br>GGCAGACTTTCTTGATATTACTCTGTCTTTCCCCAGGGCGACACCAGCACCCTCGGACACC<br>TGTGACCTCTTCCTTCTCTGCAGGAGGAGAGCCAGTGGCAACCCTACGCTGATAAGATTAG<br>TCTGAAAGGCCGATTATGGTGTAATTTCCTATGCTGAAACTTTGTAGATTTAAAAAAAAAG<br>GGGGGGGAGGGGTGTTTCATTTTCCAATTGGTCTGATTTCACAGGAAAATTTACCTGTTTC<br>TCTTTTTTCTCCTGGAAGAGAGGTGCTGATTGGCCCAGAATAACTGACAATCTGGTGTAAT<br>AAAATTTTCCAATGTAAACTCATTTTCCCTTGGTTTCAGCAACTTTAACTCTATATATAGAG<br>AGACCTCTGCCAGCATTGCATTGTTAGCATCTCTTGATAAACTTAATTGTCTCTCGTCACTG<br>ACGGCACAGAGCTATTGATGGGTCTCAACCCCCAGCTAGTTGTCATCCTGCTCTTCTTTCTC<br>GAATGTACCAGGAGCCATATCCACGGATGCGACAAAAATCACTTGAGAGAGATCATCGGC<br>ATTTTGAACGAGGTCACAGGAGAAGGGGTAAGTACCTATCTGGCACCATCTCTCCAGATA<br>CCCAGGTGATACTGCTGGGGCGATTCTAGGCTTGGAGAGCTGAGTTGCTAGAGAGGTGGA<br>CGGACGGCAGGTGGCTGAGGCAGGACTAGGGACAAAGCTCAAGGGATCCAAGCTGGGCC<br>CTGGTAATATAACTTCGTATAGCATACATTATACGAAGTTATGGAGAGGAGGAAACAACC<br>TACTTGAGGGCAGACCCTGCTACTTTTGGAGGCAGAGGGAGTCTCCCGGGTGGGGGGTGG<br>GAGGTGTAGCGATGCTTTCTCTGTCC |

**Supplementary Table 3.** List of oligonucleotides used for ssDNA donor template generation by asymmetric PCR and PCR analyses of targeted loci.

Oligonucleotides used for ssDNA donor template generation by asymmetric PCR

| **Gene name** | **Oligonucleotide ID name** | **Nucleotide sequence (5'-3')** |
|---|---|---|
| *S100A8* | A8_ssD | GCCATTTTCTGGGTTTAGTTATCTGAGTGGTACAGGCCAGGTCAAAGCTTATTCTTGGGTTAAAAAC |
| | A8_ssR | ACTATCTGTGGATAAGTATTGGCTAACTGCAGAAAGCAACTTGGAGGGCTAGAAATCCCAGCACCATTAGAAAAC |
| *Trek1* | TREK1_ssD | AGAGACAAAGGATTATCTACCATACCTGTGTTCTCTGTGCGTCTGGGACATCCCGAACAGGTGTAGGTGGGAT |
| | TREK1_ssR | CAAGTTTGCATGTGCACACACACACAGAGAAAATAATTAGGACAAAATACAACTTCAA |
| *Inf2* | INF2_ssD | CCCTGAGGCCAGAGACTTCCTCAGAAGCTCCAGGCATCCTGGGCTGCTACCAAGTTGCCCTTGCCTGACTTCGGAT |
| | INF2_ssR | GACTCTGGGATGCTACAGGAACTCCTTCCAGTCTGTGGCACCCCAACACCCCAAGTGCTCAG |
| *Trpc6* | TRPC6_ssD | GAGAGCCACAGCAGCAGAATCACCTAATCATTGGAGAGAGAGAAATGAACCTGAGTTGTAATTTGCATATATGGATC |
| | TRPC6_ssR | TGTCATGTAATCTGACTCTAAAATCATTACATATAATTATTTTATATTGGGAAGGTTGATTTATGCTAGTCTT |
| *IL4_5'LoxP* | IL4_LoxP1D | CAGCCATTTCTCAGGCTTCTGTCTAAGGTAGGAAAAATCTTCAACCTAGCCCAGAACCTCGGATCCAAGCTGGGCCCTGGTAAT |
| | IL4_LoxP1R | CTTTGGAGACAGAGAGGTGACATGGAGTGGGGCAGGCAGGACCATGGAATGAGGCTTTACCTCGAGCCCTCAAGTAGGTTGTTTCC |
| *IL4_flox* | IL4_ssD | AGCCATTTCTCAGGCTTCTGTCTAAGGTAGGAAAAATCTTCAACCTAGCCCAGAACCTTTATATAGGTAAAGCCTCATTCC |
| | IL4_ssR | GGACAGAGAAAGCATCGCTACACCTCCCACCCCCCACCCGGGAGACTCCCTCTGCCTCCAAAAGTAGCAGGGTCTGC |

Oligonucleotides used in PCR analysis.

| **Oligonucleotide names** | **Oligonucleotide sequence** |
|---|---|
| *S100A8*_d1 | CAGGTGAGGAGGTGCTATGTTC |

| | |
|---|---|
| *S100A8*_d3 | GCAGGAAGTGTTTAGTGTGGAG |
| *S100A8*_d4 | GCGTAGAGCCTTCTAGCAGTGTC |
| *S100A8*_d7 | CTGCAGTTAGCCAATACTTATCCACA |
| *S100A8*_r3 | GGGGGGGGTGAGTTCAGAC |
| *S100A8*_r4 | CAAGTTTTCGATATTTATATTCTGTCAAG |
| *S100A8*_r7 | GCCTGTACCACTCAGATAACTAAACC |
| *LoxP*_Ad1 | AAGCTGGGCCCTGGTAAT |
| *LoxP*_Ar1 | GCCCTCAAGTAGGTTGTTTCC |
| *LoxP*_Ad2 | CCAGGTTAGGATCGCATAGG |
| *LoxP*_Ar2 | CGACTACGCGCATTAAGGA |
| *IL4*_SD1 | CTTCTGTCTAAGGTAGGAAAAATCTTCA |
| *IL4*_SR1 | GAGTGGGGCAGGCAGGACC |
| *IL4*_SD1r | AGATTTTTCCTACCTTAGACAGAAGCC |
| *IL4*_SR1d | TCCTGCCTGCCCCACTCCA |
| *INF2*_d2 | CCCTGAGGCCAGAGACTTCC |
| *INF2*_r1 | GACTCTGGGATGCTACAGGAACTC |
| *Trek*_d1 | CCTGTGTTCTCTGTGCGTCTG |
| *Trek*_r1 | CATTGCCTTTCACGGACCTC |
| *Trek*_d2n | ACTGAAAATTATGTACAAAAAAACAGAA |
| *Trek*_r2 | GCACACACACACAGAGAAAATAAT |
| *Trpc*_d1 | CAGCAGCAGAATCACCTAATCA |
| *Trpc*_r2 | TGTCATGTAATCTGACTCTAAAATCA |
| *Ccnd2*_1D3 | ATGCAAAGGCTGATGCTCTGAC |
| *Ccnd2*_2R3 | CCCCAGCTAACCTATCTATAAACACTC |
| dd5LoxP_Dir | GAAGTTATGGCTGCGGAAAC |
| dd5LoxP_Rev | CAAGGCCTTCTCCAGTTCAG |
